# Modeling individual variability in habitat selection and movement using integrated step-selection analyses

**DOI:** 10.1101/2023.07.31.551213

**Authors:** Nilanjan Chatterjee, David Wolfson, Dongmin Kim, Juliana Velez, Smith Freeman, Nathan M. Bacheler, Kyle Shertzer, J. Christopher Taylor, John Fieberg

**Affiliations:** Department of Fisheries, Wildlife, and Conservation Biology, University of Minnesota; Southeast Fisheries Science Center, National Marine Fisheries Service, Beaufort, NC 28516, USA; National Centers for Coastal Ocean Science, National Ocean Service, National Oceanic and Atmospheric Administration, Beaufort, NC 28516, U.S.

**Keywords:** Animal movement, Individual variability, Integrated step-selection analysis, *Lutjanus campechanus*, Mixed-effect, Random effect, Red snapper, Step-selection function

## Abstract

*1. Integrated step-selection analyses* (ISSAs) are frequently used to study habitat selection using animal movement data. Methods for incorporating random effects in ISSAs have been developed, making it possible to quantify variability among animals in their space-use patterns. Although it is possible to model variability in both habitat selection and movement parameters, applications to date have focused on the former despite the widely acknowledged and important role that movement plays in determining ecological processes from the individual to ecosystem level. One potential explanation for this omission is the absence of readily-available software or examples demonstrating methods for estimating movement parameters in ISSAs with random effects.

2. We demonstrated methods for characterizing among-individual variability in both movement and habitat-selection parameters using a simulated data set and by fitting two models to an acoustic telemetry data set containing locations of 35 red snapper (*Lutjanus campechanus*). Movement kernels were assumed to depend on either the type of benthic reef habitat in which the fish was located (model 1) or the distance between the fish’s current location and nearest edge habitat (model 2). In both models, we also quantified habitat selection for different benthic habitat classes and distance to edge habitat, and we allowed for individual variability in movement and habitat-selection parameters using random effects.

3. The simulation example highlights the benefits of a mixed effects specification, namely we can increase precision when estimating individual-specific movement parameters by borrowing information across like individuals. In our applied example, we found substantial among-individual variability in both habitat selection and movement parameters. Nonetheless, most red snapper selected for hardbottom habitat and for locations nearer to edge habitat. They also moved less when in hardbottom habitat. Turn angles were frequently near ±*π*, but were more dispersed when fish were far away from edge habitat.

4. We provide code templates and functions for quantifying variability in movement and habitat-selection parameters when implementing ISSAs with random effects. In doing so, we hope to encourage ecologists conducting ISSAs to take full advantage of their ability to model among-individual variability in both habitat-selection and movement patterns.

## 1 Introduction

Our understanding of how individuals move and select for different habitat features has been enhanced by technological advances, including smaller and better tracking devices (Kays et al. 2015), widespread availability of remote sensing data products (Dodge et al. 2013; Neumann et al. 2015), and new statistical methods and software for modeling animal movement (Hooten et al. 2017; Joo et al. 2020). By tracking individuals in the context of dynamic environmental conditions, we can identify drivers of movement and predict how climate and land-use change might influence both individual-animal fitness and population-level characteristics (Cagnacci et al. 2010).

*Integrated step-selection analyses* (ISSAs) have emerged as one of the most popular frameworks for analyzing animal telemetry data (Avgar et al. 2016; Fieberg et al. 2021; Northrup et al. 2022). Similar to species distribution models and resource-selection functions, ISSAs infer the importance of environmental covariates by comparing their distribution at used versus available locations. Integrated step-selection analyses offer several advantages over the former approaches, however, by: 1) allowing for temporally dynamic covariates that drive species distribution patterns to be included in the model; 2) using an individual’s movements to determine available habitat; and 3) relaxing the assumption of independence among locations to an assumption of independent “steps” or movements between sequential locations. Another advantage of ISSAs is the ability to model both habitat selection, using covariates at the end of the movement step, and habitat-dependent movement, by allowing an animal’s step lengths (distances between sequential locations) and turn angles (changes in direction) to depend on habitat at the start of the movement step (Avgar et al. 2016; Fieberg et al. 2021). This strength has been widely recognized, and ISSAs have been used to explore how individuals alter their step lengths and turn angles in response to linear features such as roads and seismic lines (Prokopenko, Boyce, and Avgar 2017; Scrafford et al. 2018; Dickie et al. 2020).

Individuals from the same species often show substantial variation in their space-use patterns, and it is appealing to model this variation using random effects (Duchesne, Fortin, and Courbin 2010). However, random effects can be computationally challenging to include within the conditional logistic regression framework typically used to estimate parameters in ISSAs (Duchesne, Fortin, and Courbin 2010; Craiu et al. 2011). This led Craiu et al. (2011) to develop a two-step approach for fitting mixed conditional logistic regression models in this context. More recently, Muff, Signer, and Fieberg (2020) developed an approach for incorporating random effects in ISSAs by exploiting the equivalence between Poisson regression with stratum-specific intercepts and conditional logistic regression. This approach has quickly become commonplace when modeling habitat selection using data sets that include many individuals, and allows for both individual- and population-level inference. For example, one can estimate coefficients for a typical individual by setting all random effects to 0 (Fieberg et al. 2009, 2010), estimate variance parameters that capture among-individual variability in the habitat-selection coefficients (Muff, Signer, and Fieberg 2020), and explore functional responses in habitat selection by considering how individual-specific coefficients vary with habitat availability or other landscape metrics (Matthiopoulos et al. 2011; Aarts et al. 2013). A nice exemplification is Jones et al. (2020), who used random effects to explore whether variation among individual spotted owls (*Strix occidentalis*) in their habitat-selection parameters was related to patch size and configuration in a post-fire landscape.

Individuals also frequently exhibit variation in the extent of their movements, which can have implications for survival and fitness. For example, Fraser et al. (2001) found that Trinidad killifish (*Rivulus hartii*) identified in the lab as being more bold also moved more when released into their native streams; killifish movements were also positively correlated with individual growth in areas of the stream where predators were present. Although many studies have explored individual variation in habitat-selection parameters, little to no attention has been given to studying variation in movement parameters when using ISSAs with random effects (hereafter “mixed ISSAs”). One likely reason for this lack of attention is that the estimation of movement parameters in ISSAs is typically done using a complicated two-step process (Avgar et al. 2016; Fieberg et al. 2021). Tentative movement parameters are first estimated using observed step lengths and turn angles, ignoring the effect of habitat selection on observed movements, and then movement parameters are updated using additional coefficients from the fitted conditional logistic regression model (Figure 1). This approach is analogous to using importance sampling to approximate the likelihood of the data (Michelot et al. 2023). The amt package (Signer, Fieberg, and Avgar 2019) provides functions for implementing this two-step approach, but they only work when fitting models to individual animals. Furthermore, it is not immediately clear how this process should be implemented when the goal is to estimate individual-specific movement parameters.

**Figure 1:**
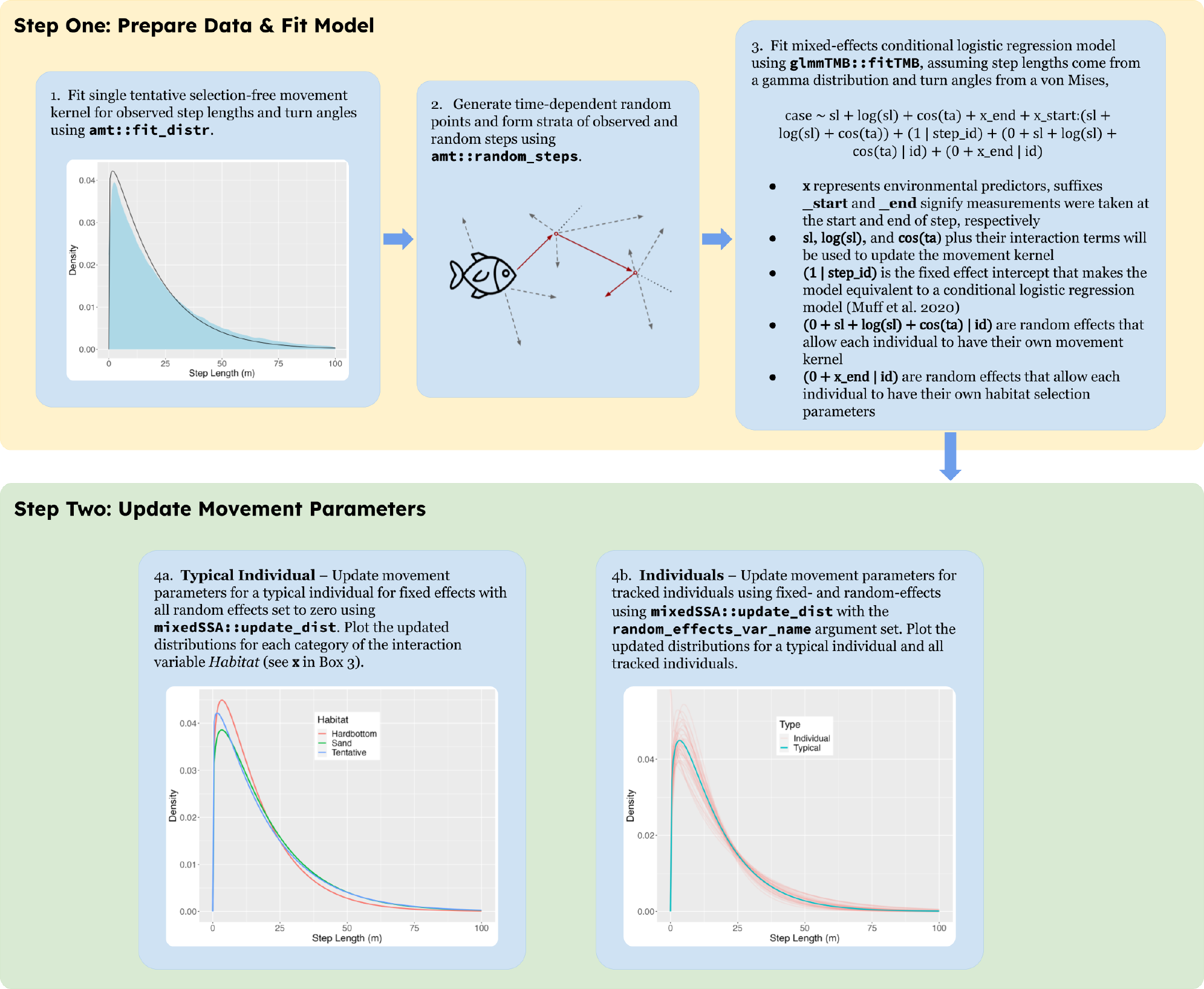
Schematic diagram of proposed workflow for estimating individual-specific movement parameters from a mixed-effects integrated step-selection analysis. Although we focus here on visualizing step-length distributions in different environments and across different individuals in the population, the same process can be used to visualize variation in turn-angle distributions. For worked examples, see Appendix 1.

Here, we demonstrate how mixed ISSAs can be used to study among-individual variation in both movement and habitat-selection parameters using a simulation example and by analyzing an acoustic telemetry data set of 35 red snapper (*Lutjanus campechanus*). We show how one can visualize individual variation in movement and habitat-selection patterns, and provide functions within an associated R package (mixedSSA) to facilitate the application of these methods to other data sets.

## 2 Integrated Step-Selection Analysis (ISSA)

Integrated step-selection analyses assume that animal space use is captured by the product of two functions, a movement-free habitat-selection function, *w*(*·*), that represents habitat preferences and a selection-free movement kernel, *ϕ*(*·*), that quantifies how the animal would move in the absence of habitat selection (Avgar et al. 2016; Fieberg et al. 2021):

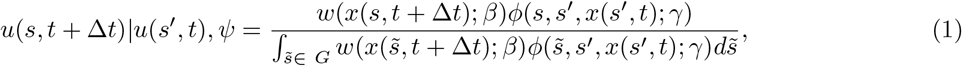

where *u*(*s, t* + Δ*t*) *u*(*s*^*′*^, *t*), *ψ* represents the conditional probability of an animal being present at location *s* at time *t* + Δ*t*, given that it was at location *s*^*′*^ at time *t* and arrived there via the bearing *ψ*; *x*(*s*^*′*^, *t*) and *x*(*s, t* + Δ*t*) are habitat covariates measured at the start and end of the movement step, respectively; and *β* and *γ* are sets of parameters that quantify habitat selection and movement tendencies, respectively. Lastly, note that the denominator in equation (1) integrates over the geographic area, *G*, that represents the area in which an animal could have potentially moved based on its prior observed movements and ensures that the right-hand side is a proper probability distribution; 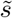 is a dummy variable for integration.

## 3 Example Application to Simulated Data

To demonstrate the application of mixed ISSAs that allow for individual variability in movement parameters (Figure 1), we simulate trajectories following equation (1) for a population of individuals with individual-specific, known parameter values. We also compare the estimates from the mixed ISSA to those obtained by fitting a separate model to data from each individual. Doing so highlights the benefits of using random effects; namely, the ability to borrow information across like individuals, which leads to improved estimates of individual-specific parameters (Gelman and Hill 2006; Harrison et al. 2018).

### 3.1 Simulated Movements

We generated a set of tracking data for 30 individuals. To do so, we created a landscape with a single categorical variable representing two habitat types (A and B). We then simulated movements in which each individual had its own set of movement and habitat selection parameters, and in which movements were more directed, on average, when individuals were in habitat of type B.

Let *sl*_*ik*_ and *θ*_*ik*_ represent step lengths and turn angles, respectively, for individual *i* in habitat class *k* (*k* = 1, 2 for habitats A and B). The *sl*_*i*1_ follow a gamma distribution with individual-specific shape and scale parameters, *α*_*i*1_ and *ν*_*i*1_ respectively, and *θ*_*i*1_ follow a von Mises distribution with mean 0 and with individual-specific concentration parameters, *κ*_*i*1_:

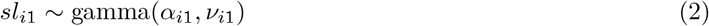

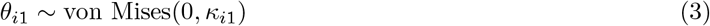

We assumed the parameters (*α*_*i*1_, *ν*_*i*1_, *κ*_*i*1_) for each individual followed a multivariate Normal distribution:

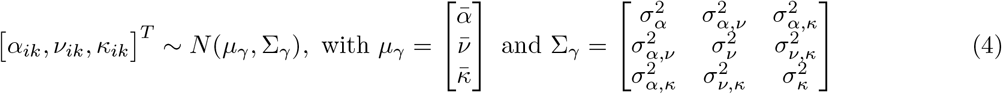

The diagonal elements of Σ_*γ*_ quantify among-individual variability in the movement parameters, and the off-diagonal elements quantify how these parameters covary; non-zero covariances allow individuals that tend to move more to also exhibit more directed movements (or, vice versa depending on the sign of the covariance parameters). We assumed that step lengths and turn angles for movements starting in habitat B also followed individual-specific gamma and von Mises distributions, respectively. The parameters of these distributions were shifted relative to parameters in habitat A using an additive model:

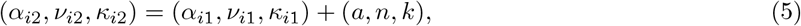

where (*a, n, k*) are constants. Thus, we allowed each type of movement parameter (*α, ν*, and *κ*) to vary by individual, while forcing *differences* in parameter values for the two habitat classes to be consistent across individuals. For example, *α*_*i*2_−*α*_*i*1_ = *a* for all individuals. Lastly, we assumed *w*(*x*_*t*+1_) = *exp*(0.5) when individuals were in habitat B and *w*(*x*_*t*+1_) = *exp*(0) when in habitat A. Thus, all individuals preferred habitat B to habitat A.

We generated random parameters, (*α*_*i*1_, *ν*_*i*1_, *κ*_*i*1_), for each individual using the `mvrnorm` function in the `MASS R` package (Venables and Ripley 2002). We then generated random trajectories consisting of between 1633 to 2627 locations using the `redistribution_kernel` and `simulate_path` functions in the `amt` package of R (Signer, Fieberg, and Avgar 2019; Signer et al. 2023). We set the parameter values used to simulate the trajectories as follows:

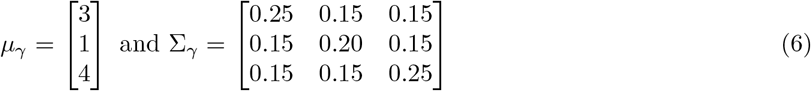

Lastly, we set (*a, n, k*) = (0.05, 0.05, 0.05).

### 3.2 Parameter Estimation

#### 3.2.1 Mixed ISSA

We used the following workflow to estimate the parameters in them mixed ISSA (Figure 1):

1. We fit a single tentative selection-free movement kernel, 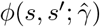, to the observed step-lengths and turn angles pooled across all individuals. Here, 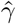 consisted of a single shape (*α*_0_), scale (*ν*_0_), and concentration parameter (*κ*_0_). We set the mean of the von Mises distribution to 0.
2. Using `amt’s random_steps` function, we generated time-dependent random locations by simulating 10 potential movements from each previously observed location using the tentative selection-free movement kernel, 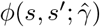from [1] above.
3. We formed strata by matching each observed step with its set of random steps. We then used the `glmmTMB` function in the `glmmTMB` package (Magnusson et al. 2017) to fit the equivalent of a mixedeffects conditional logistic regression model using these strata and the “Poisson trick” from Muff, Signer, and Fieberg (2020). We included the habitat class measured at the end of the movement step to quantify relative selection for the two habitat types. As described in Avgar et al. (2016) and Appendix C of Fieberg et al. (2021), we can obtain updated estimates of movement parameters that account for habitat selection if we include specific movement characteristics in our conditional logistic regression model; these characteristics depend on the assumed distributions used to model step lengths and turn angles. For our model specification using the gamma and von Mises distributions, we need to include step length, log(step length) and cos(turn angle). Further, to allow the movement parameters to depend on habitat class, we need to include interactions between these movement characteristics and the habitat class at the start of the movement step. Lastly, we included random effects for step length, log(step length), and cos(turn angle) to allow each individual to have its own set of movement parameters.
4. We updated the movement parameters in *ϕ*(*s, s*^*′*^; *γ*) using the regression coefficients associated with movement characteristics (step length, log(step length), and cosine(turn angle)) and their interactions with habitat class as outlined in Avgar et al. (2016) and Appendix C of Fieberg et al. (2021). Let *β*_*sl*_, *β*_log(*sl*)_, and *β*_*cos*(*θ*)_ represent the fixed-effects coefficients associated with step length, log(step length) and cos(turn angle), respectively. Further, we let 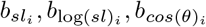 represent individualspecific random effects associated with these variables. We estimated updated movement parameters for the reference class (habitat A) for each individual, *i*, using:

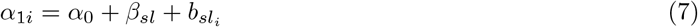

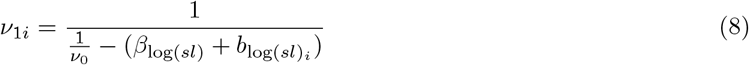

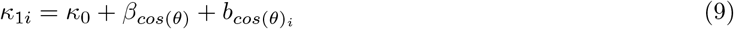

Similarly, we could estimate updated parameters for habitat class B by including the coefficients for the interaction terms in the above equations (see e.g. Appendix B of Fieberg et al. 2021 or Appendix 1 for a similar application to the snapper data).

#### 3.2.2 Individual Models

We followed a similar workflow when fitting models to individuals, except that we estimated a separate tentative selection-free movement kernel using each individual’s data in step 1, and we used the `fit_issf` function in the amt package when fitting models to each individual’s data rather than `glmmTMB` in step 3; the `fit_issf` function serves as a wrapper function for the `clogit` function in the survival package (Therneau 2023).

## 4 Application of Mixed-ISSAs to Red Snapper

Red snapper are an economically and ecologically important species distributed in the Gulf of Mexico and along the eastern coasts of North America, Central America, and northern South America. They are a reef-associated, demersal, gonochoristic fish species that can live 50+ years and matures around two years old (Wilson and Nieland 2001; White and Palmer 2004). The size of individuals can exceed 1000 mm and 12 kg. They are generalist predators, with adults feeding primarily on benthic invertebrates and other reef-associated fishes (Szedlmayer and Lee 2004; Wells, Cowan Jr, and Fry 2008). In the United States, red snapper are targeted by both recreational and commercial sectors, and these fisheries are managed in the Gulf of Mexico by the Gulf of Mexico Fishery Management Council and in the Atlantic Ocean by the South Atlantic Fishery Management Council. Previous studies of red snapper have found an affinity for areas of high to moderate relief near edges of hardbottom habitat (Williams-Grove and Szedlmayer 2016, 2017; Bacheler et al. 2021; Bohaboy, Cass-Calay, and Patterson III 2022).

### 4.1 Study Area

Our study site comprises reef habitat in approximately 37 m deep water off the coast of North Carolina, USA, between Cape Hatteras and Cape Lookout (Figure 2). The seafloor of the study site is composed of a patchy mix of rocky pavement and ledges in a matrix of sand (Figure 2). We used a high-resolution bathymetric map for classification of benthic habitat into either sand or hardbottom.

**Figure 2:**
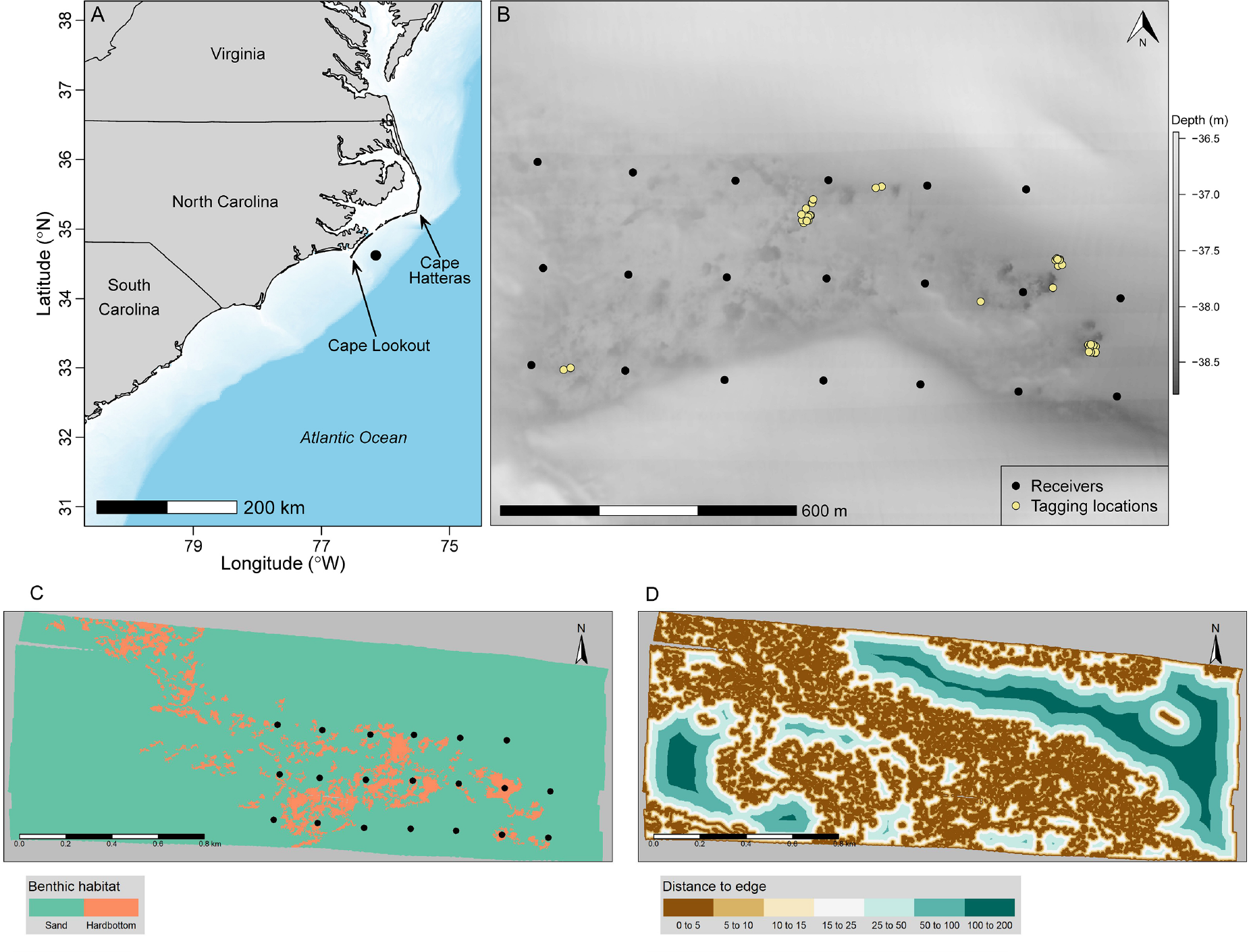
(A) Broad-scale location of study (filled black circle) on the continental shelf between Cape Lookout and Cape Hatteras, North Carolina, USA, in 2019. Water depth is shown in blue: the lightest blue indicates water less than 5 m deep, while the darkest blue is greater than 100 m deep. (B) Close-up view of the study area where a Vemco positioning system was used to estimate fine-scale positions of red snapper (*Lutjanus campechanus*). Background shows a multibeam sonar map that is scaled to depth (right y-axis), submersible receivers are represented by filled black circles, and tagging and release locations are represented by light yellow points. Environmental variables used in the habitat-selection modelling (C) benthic habitat class (sand or hardbottom) with the submersible receivers represented by filled black circles; and (D) distance-to-edge habitat.

### 4.2 Data Collection and Processing

We caught red snapper from May through September, 2019 using hook-and-line methods or baited fish traps and tagged them with 13g externally-attached Vemco V13P-1x transmitters that had an average battery life of 613 days and a pulse interval of 130-230 seconds. Locations were collected from May 2019 to December 2019 using a 400 x 1200 m regularly spaced grid array of 20 VR2AR acoustic receivers placed on the sea floor (Figure 1, Bacheler et al. 2021). We used an additional V13P-1x reference transmitter to quantify the accuracy of data collection between transmitters and acoustic receivers by deploying it at a known location and quantifying the horizontal error of all estimated positions over the duration of the study.

We thinned the telemetry data to equally spaced intervals of 6±1 minutes using the `track_resample` function in the `amt R` package (Signer, Fieberg, and Avgar 2019). We then converted the location data to a set of steps representing movements between sequential observations and dropped any steps that had a step length > 100 meters and a turn angle > 170 degrees, assuming these steps likely resulted from erroneous locations with large measurement error (Bjørneraas et al. 2010). Lastly, we created a *distance-to-edge* habitat covariate measuring the nearest distance to habitat separating sand and hardbottom habitat classes (Figure 2).

### 4.3 Mixed ISSAs

We hypothesized that red snapper would 1) prefer hard bottom habitat near the edge of reef-sand habitat transition; and 2) move in a more directed manner (longer step-lengths with turn angles near 0) when in sandy habitats or far from the edge of reef-sand habitat. To test these assumptions, we used two different mixed ISSAs. In the first, we assumed step lengths and turn angles depended on the categorical benthic habitat class variable measured at the start of the movement step, similar to our simulation example. In the second mixed ISSA, we modeled the distribution of step lengths and turn angles as a function of a continuous variable quantifying the distance between the current location and the nearest edge habitat. To allow the movement kernels, *ϕ*, to depend on the habitat at the start of the movement step, we included step length, log(step length), and cos(turn angle) as well as interactions between these terms and the benthic habitat class (model 1) or distance to edge (model 2). In both models, we also included random effects for step length, log(step length), and cos(turn angle), thus allowing each individual to have their own set of movement parameters. We did not, however, include random effects for interaction terms between step length, log(step length), cos(turn angle) and habitat class (model 1) or distance to habitat edge (model 2). Although in principle, these interactions could also be modeled using random effects, this would lead to a substantially larger number of variance/covariance parameters that would need to be estimated with relatively few individuals.

For both mixed ISSAs, we modeled the habitat-selection function, *w*(.), as a log-linear function of benthic habitat class and distance-to-edge measured at the end of the movement step:

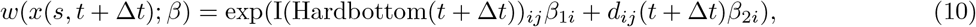

where *i* indexes the different individuals and *j* indexes the movement steps, *d*_*ij*_(*t*+Δ*t*) is the distance to edge habitat at the end of the movement step, and I(Hardbottom(*t*+Δ*t*))_*ij*_ is equal to 1 if the end of the movement step falls in the Hardbottom habitat class, and 0 otherwise. We used random effects to allow each individual to have its own set of habitat-selection parameters. We assumed the coefficients for Hardbottom habitat varied independently from the distance-to-edge coefficients and that both habitat-selection parameters varied independently of the movement parameters.

We estimated parameters using the same workflow as outlined in Figure 1 and implemented in the application to the simulated data. We noted, however, that the peak of the snapper turn-angle distribution was centered on ±*π*. This can occur when individuals are largely stationary or moving back and forth along linear features. To allow for a more flexible specification of the von Mises distribution, we set the estimated mean of the turn-angle distributions to ±*π* and |*κ*| to *κ* when estimates of *κ* were negative and 0 otherwise. Lastly, we estimated the mean step length for a typical individual (i.e., one with all random effects set equal to 0) in sand and hardbottom habitat classes (model 1) and for quantiles of *d*_*ij*_(*t*) (model 2). We used the delta method to quantify uncertainty in these estimates.

We provide a full set of equations describing the two models along with the R code needed to implement them in Appendix 1.

## 5 Software and Data Availability

We developed an R package, mixedSSA (available at https://github.com/smthfrmn/mixedSSA) to facilitate estimation and visualization of updated movement parameters. All analyses were carried out in R 4.0 (R Core Team 2020). Data and code for reproducing all of the analyses will be archived in the Data Repository of the University of Minnesota upon acceptance.

## 6 Results

### 6.1 Simulation Example

As expected, the mixed ISSA resulted in more accurate estimates of the movement parameters than when fitting separate models to each individual. Deviations between the estimated and true parameters were smaller, on average, for the mixed ISSA (Figure 3), with Root Mean-Squared Errors (RMSEs) that were roughly half that of the individual model fits. The RMSEs for the shape, scale, and kappa parameters were 0.09, 0.035, and 0.10 for the mixed-effects model versus 0.18, 0.077, and 0.21 for the individual models.

**Figure 3:**
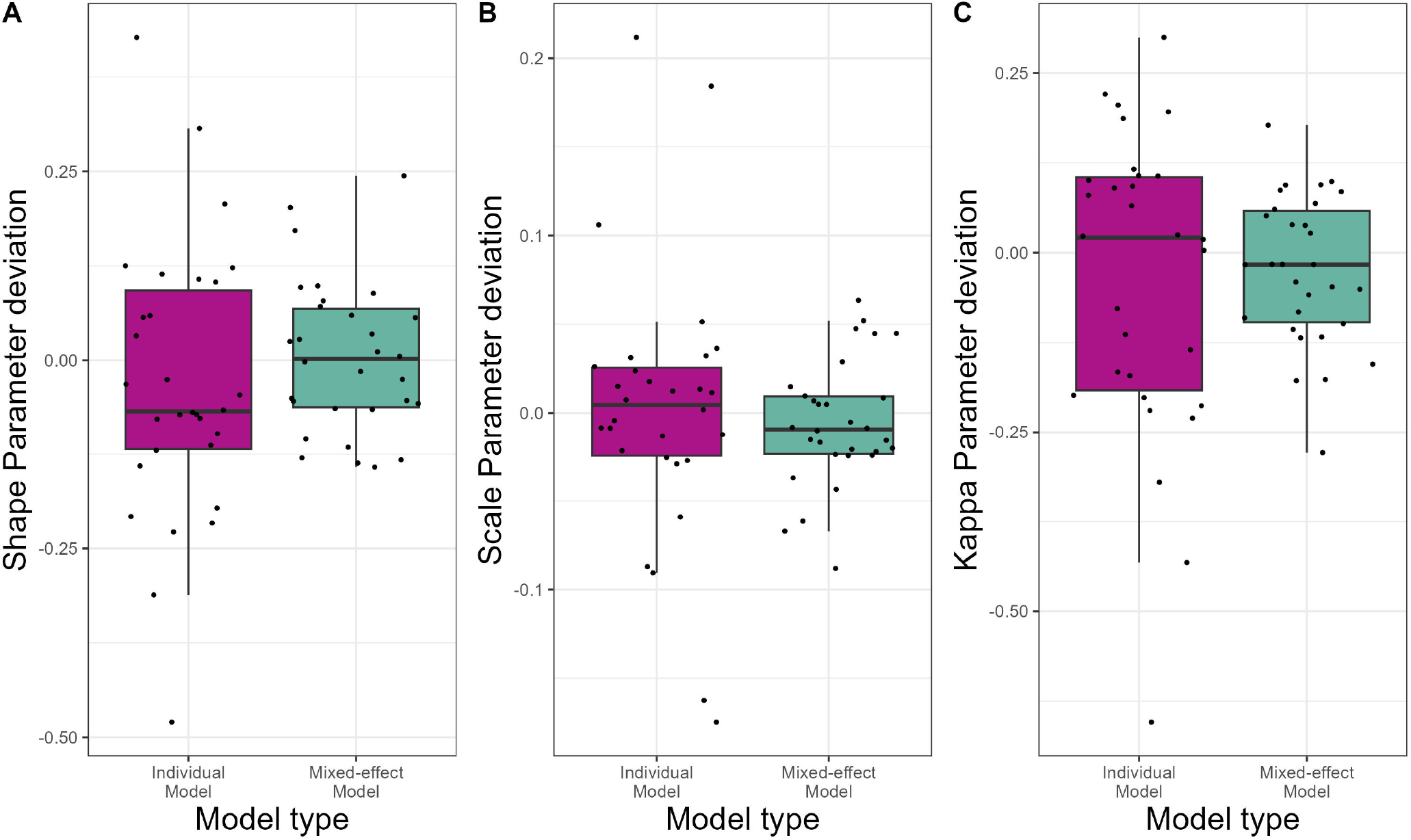
Deviations (estimated−true) in the shape (*α*), scale (*ν*), and kappa *κ* movement parameters estimated using the individual (purple) and mixed-effect ISSA (green) models (A−C) for 30 simulated individual tracks.

### 6.2 Application to Red Snapper

A total of 35 individuals were acoustically tagged in this study. The number of relocations (range 265 — 52,189) as well as the number of tracking days (range 3 — 223) were substantially different across individuals (Supplementary Table 1). Median horizontal error from 18,026 detections of the reference transmitter was 0.8m with a range of approximately 0.6 — 2.0m (Bacheler et al. 2021). We dropped a total of 13,802 observations (4% of the total) that did not satisfy the filtering criteria.

There was substantial variability in the estimated individual-specific habitat-selection parameters, with both positive and negative coefficients for each explanatory variable (Figure 4, Supplementary Figure 8). Comparing mean coefficients and their SEs, we see that the “typical individual” prefers hardbottom to sand habitat (Figure 4), though the coefficient for I(hardbottom) was negative for 3 individuals.

**Figure 4:**
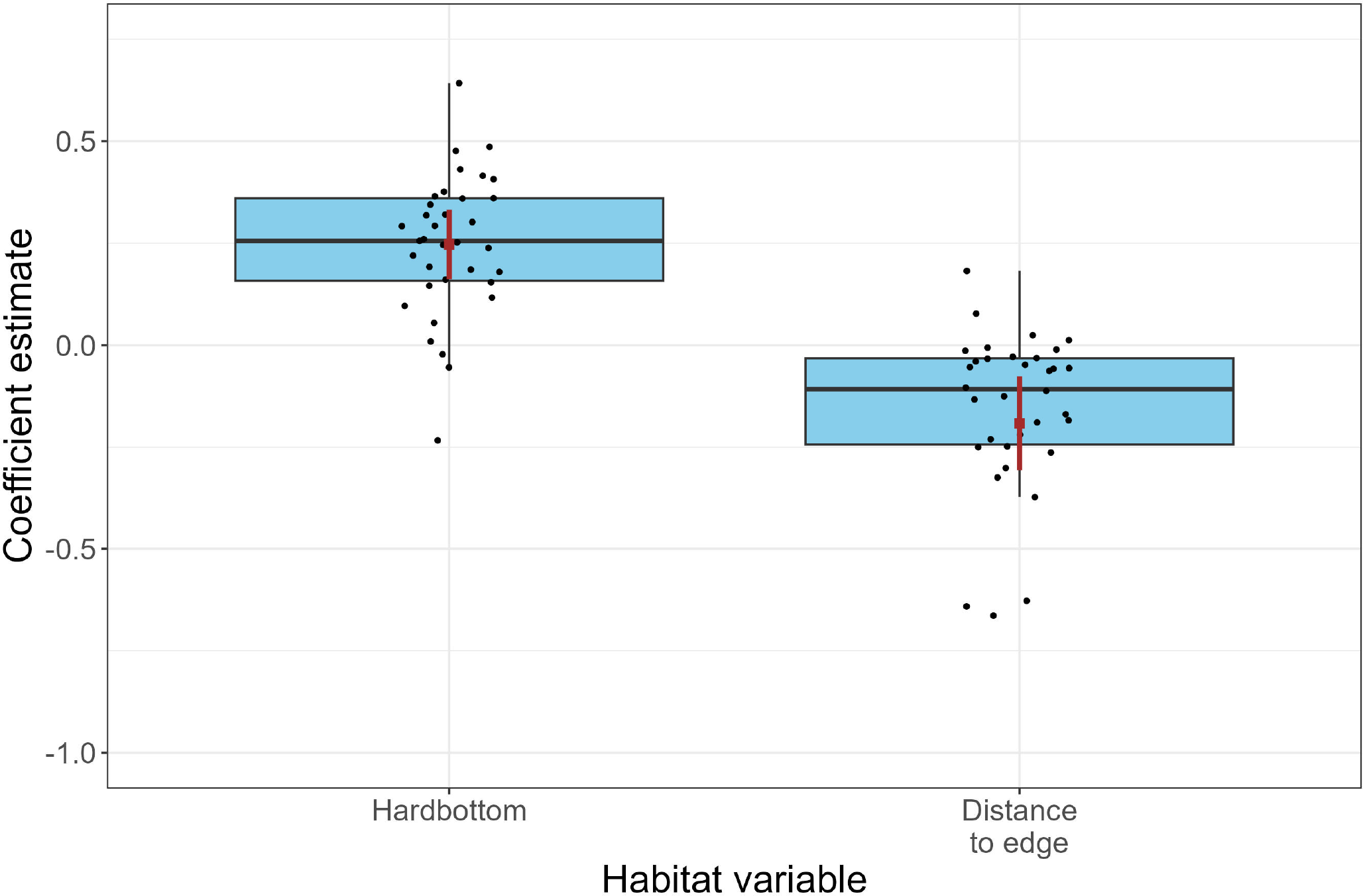
Boxplots summarizing the individual habitat-selection coefficients (black dots) associated with hardbottom habitat and the distance-to-edge covariate from Model 1. Boxes bound the 25th and 75th percentiles, solid line within the box indicates the median, and the whiskers extend to 1.5 times the interquartile range of the observations. The mean coefficient, along with a 95% confidence interval for the mean, is depicted in brown.

Evaluating movement kernels for an individual with random effects = 0 (Figure 5, Supplementary Table 3), we see that the typical snapper also tends to move less when in hardbottom habitat and as they get closer to edge habitat. The distribution of turn angles also becomes more dispersed when fish are farther from habitat edges (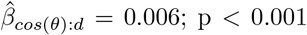Figure 5). Similar to habitat selection, there was substantial among-individual variability in both the distributions of step-lengths and turn angles (Figure 6, Supplementary Figure 9) in both models.

**Figure 5:**
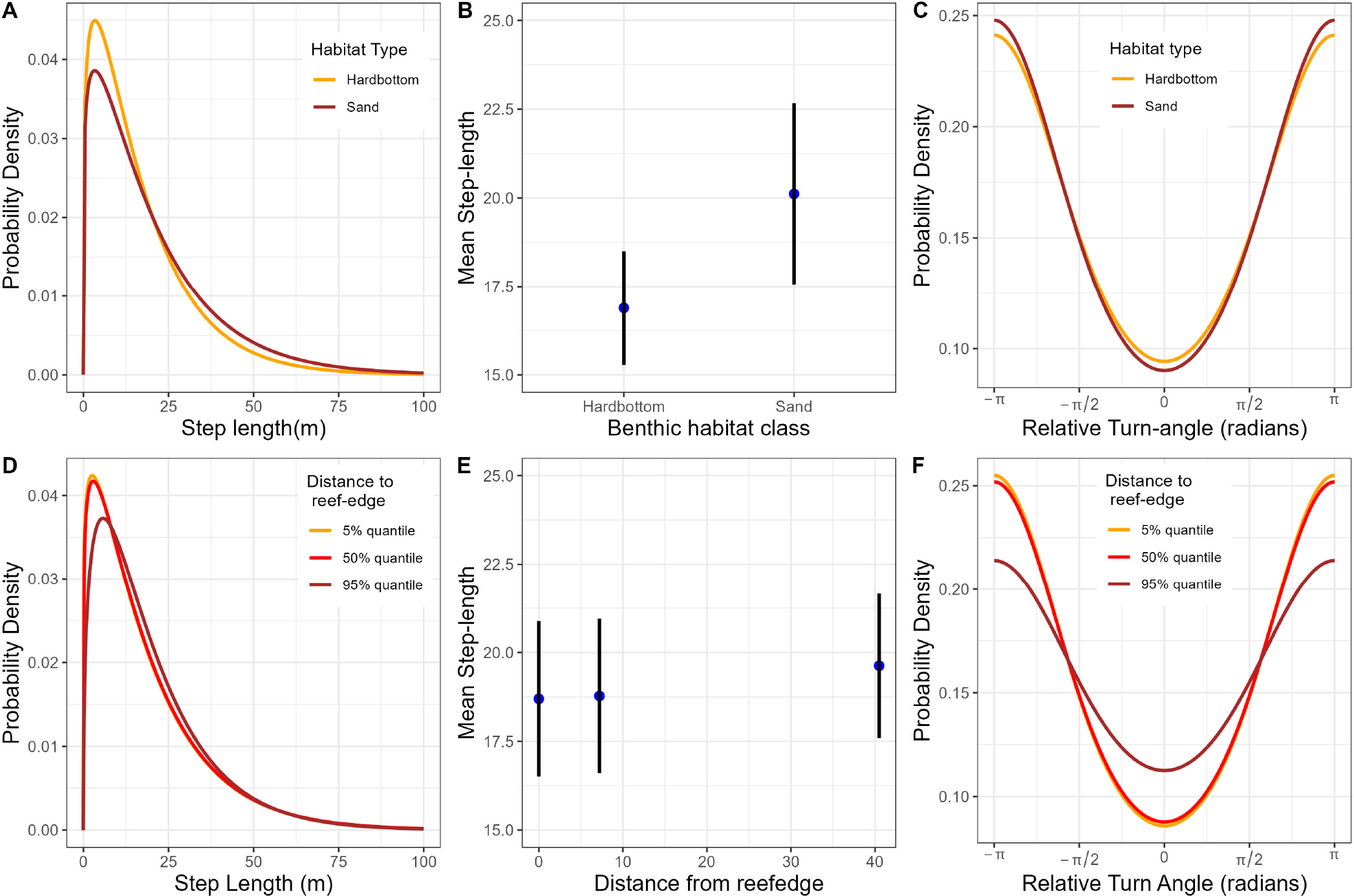
Estimated step-length and turn-angle distributions for a typical red snapper, i.e., one with all random effects equal to 0, for sand and hardbottom habitat classes (A, C) and different quantiles of distance to edge habitat (D, F) with estimates of mean step length (i.e., speed) for sand and hardbottom habitat class (B) or for different quantiles of distance to edge habitat (E).

**Figure 6:**
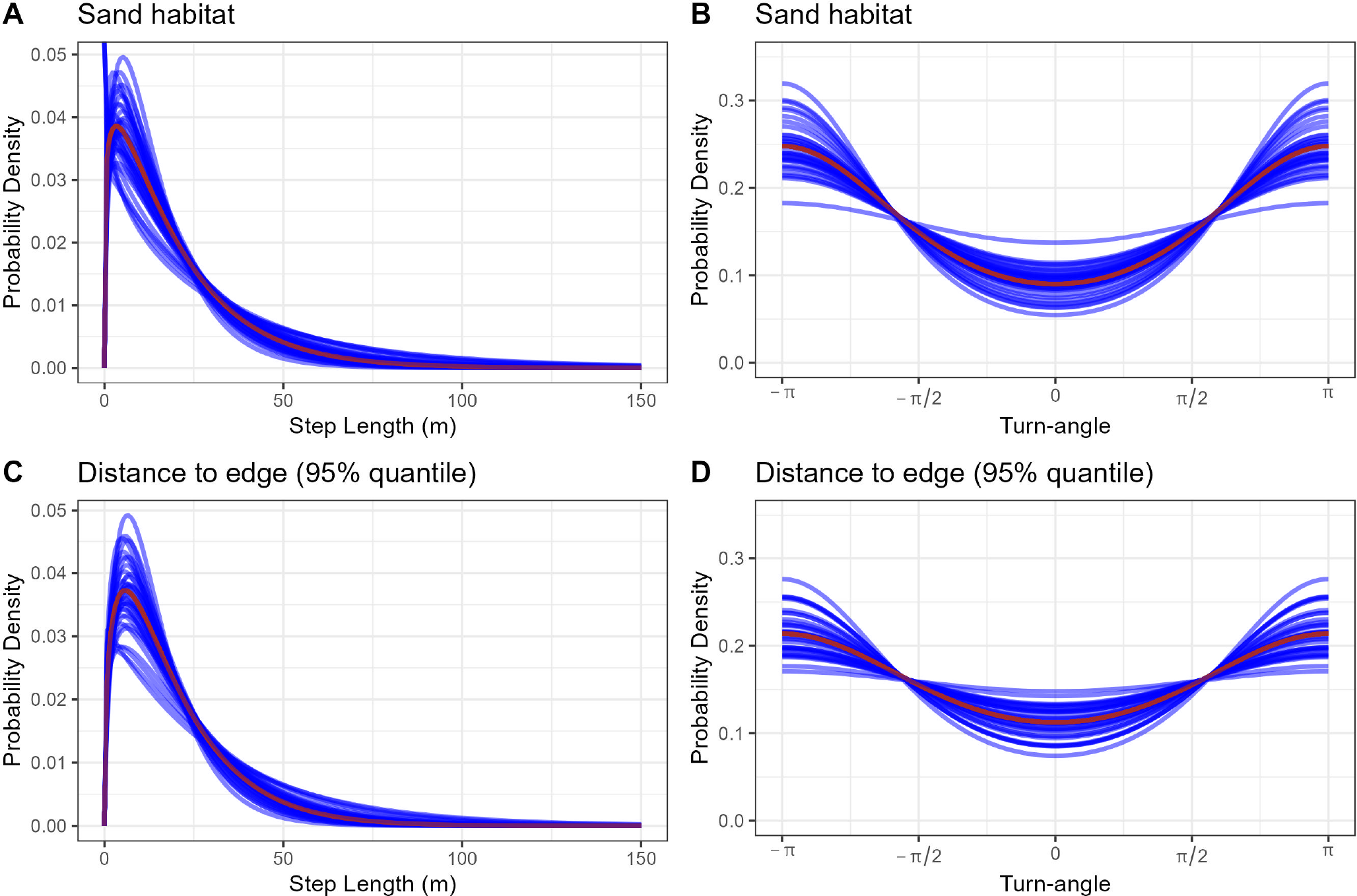
Estimated step-length and turn-angle distributions for different individuals (blue) and for a typical individual (i.e., an individual with all random effects set to 0) (red) in sandy habitat (A-B) and at 95% distance quantile from edge habitat (C-D).

**Figure 7:**
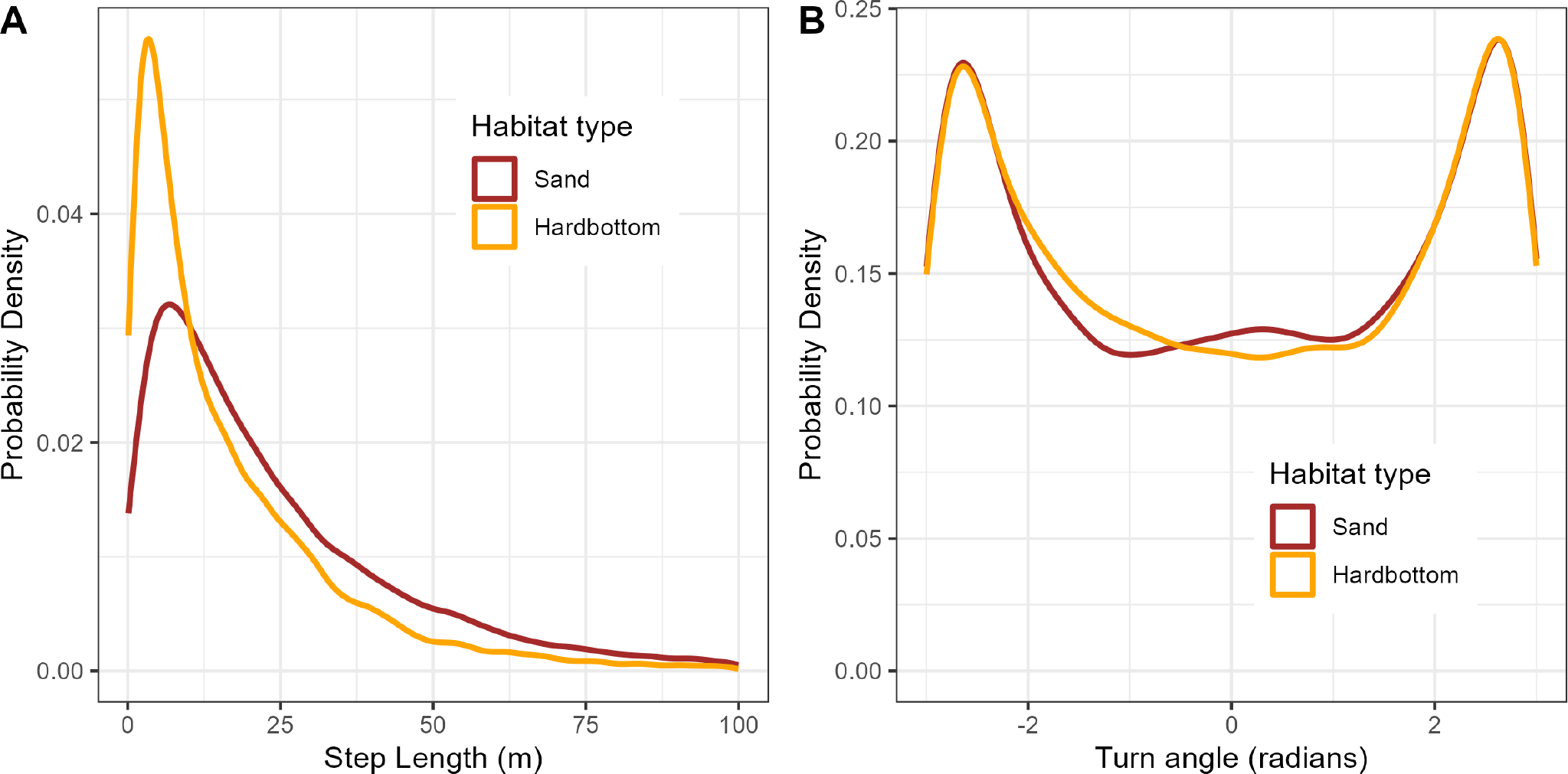
Empirical distribution of step lengths (m) and turn angles (radians) using data pooled across 35 red snappers and resampled with a fix-interval of six minutes

**Figure 8:**
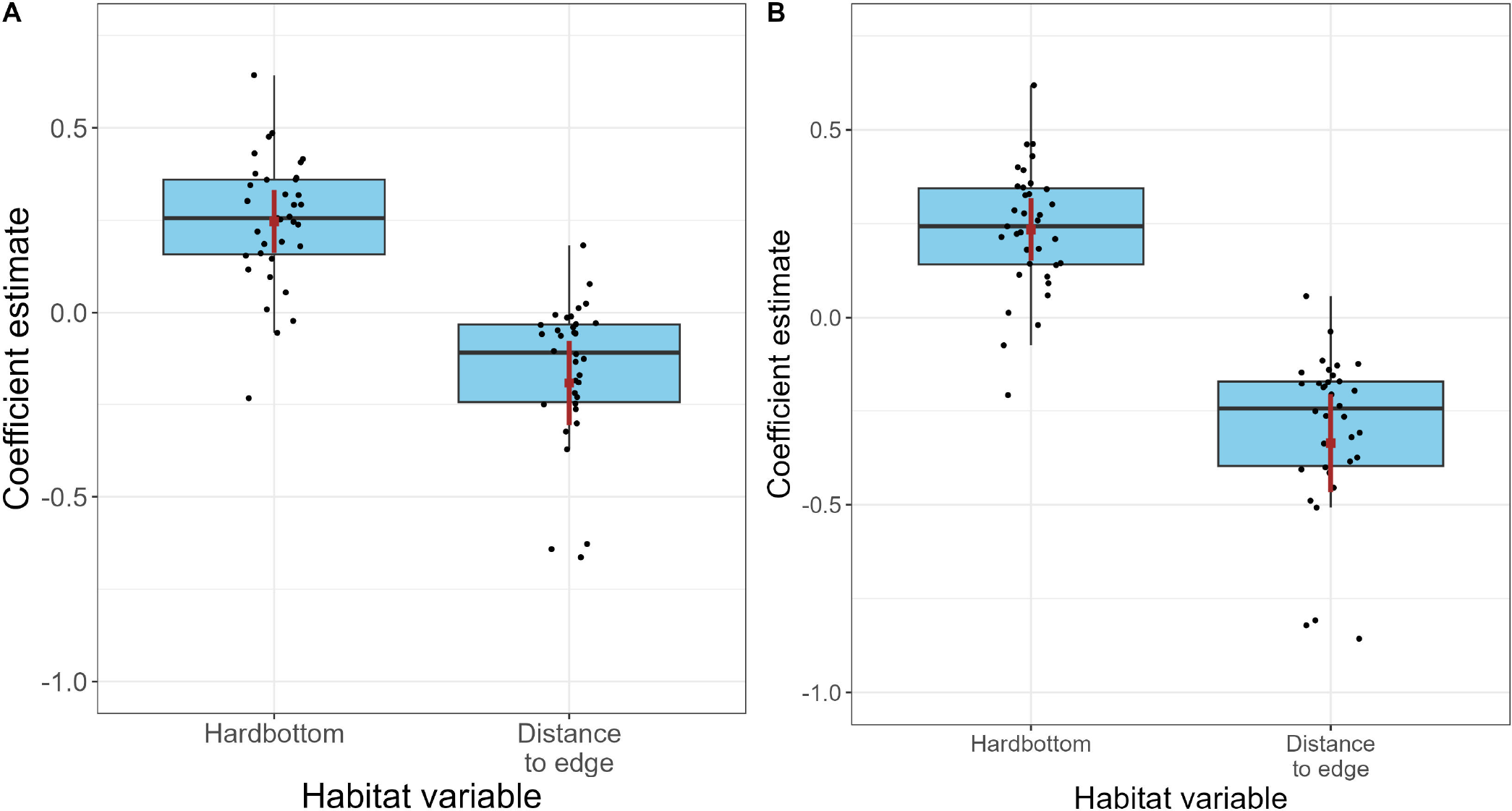
Box plot summarizing the individual habitat-selection coefficients (black dots) associated with hardbottom habitat and the distance-to-edge covariate from Model 1 (A) and Model 2(B). Boxes bound the 25th and 75th percentiles, solid line within the box indicates the median, and the whiskers extend to 1.5 times the interquartile range of the observations. The mean coefficient, along with a 95% confidence interval for the mean, is depicted in brown.

**Figure 9:**
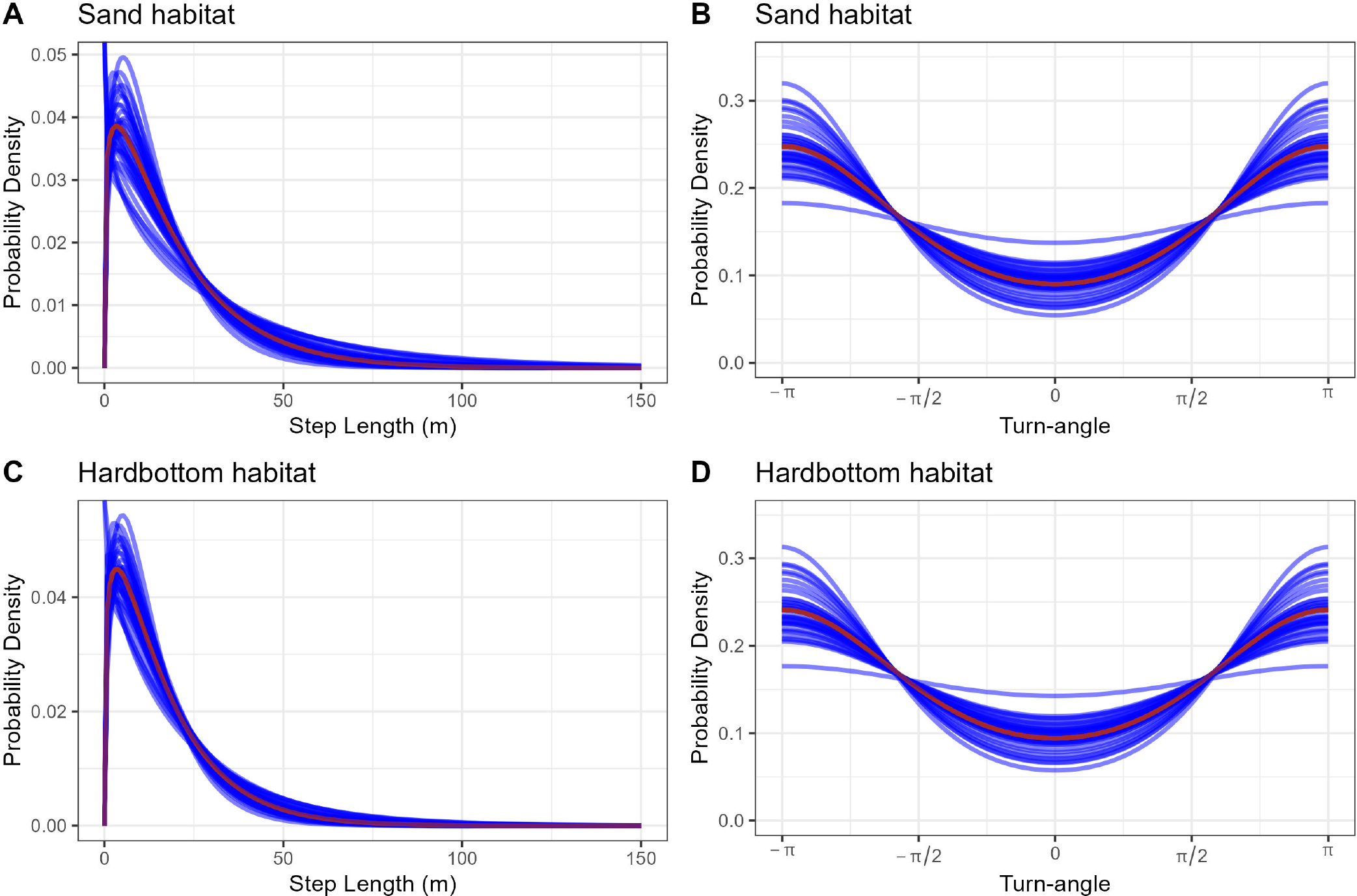
Estimated step-length and turn-angle distributions for different individuals (blue) and for a typical individual (i.e., an individual with all random effects set to 0)(red) in sand (A-B) and hardbottom habitats (C-D).

## 7 Discussion

An understanding of the causes and consequences of variability in animal movement and habitat selection is important for addressing many questions in applied and theoretical ecology (Shaw 2020). For example, behavioral ecologists are increasingly interested in studying personalities and behavioral syndromes using mixed-effect models applied to metrics of animal movement and habitat selection (Hertel et al. 2019, 2020). Consistent and quantifiable intra- and inter-individual variability in movement also drives many important ecological processes, including individual foraging rates, social interactions, disease dynamics, and spread of invasive species (Spiegel et al. 2017; Webber et al. 2020, 2023).

Our application to data from red snappers tracked on reef habitat demonstrates how mixed ISSAs can be used to study both intra- and inter-specific variation in animal movements. By including interactions between movement attributes and environmental covariates (benthic habitat class or distance to edge of hardbottom habitat), we were able to quantify how individuals altered their movements depending on local habitat features. Interestingly, we found that individuals tended to have shorter average step lengths in the most strongly selected habitat (i.e., hardbottom habitat). Turn angles also tended to be more dispersed when fish were close to edges of reef habitat. This combination of shorter step lengths and more dispersed turn angles is consistent with area restricted search patterns, suggesting that these locations may serve as important foraging areas. Similar interactions with temporal covariates could be used to study how individuals alter their movements through time (e.g., Forester, Im, and Rathouz 2009; Giroux et al. 2023).

Random effects associated with each individual allowed us to quantify among-individual variability in both habitat-selection parameters and movement kernels. As is often the case, we found substantial among-individual variability in habitat-selection parameters; we also found substantial variability in individual movement kernels. Individual variability in red snapper movement and habitat use likely has important management implications. For instance, red snapper vulnerability to capture by fishers is likely related to their movement and habitat use, which could create the potential for evolutionary effects of fishing (Claireaux, Jørgensen, and Enberg 2018). Similarly, individual variability in behavior likely affects detection probabilities of red snapper sampled by surveys, and this heterogeneity may bias abundance estimates. Indeed, abundance estimates from surveys are the most important data source on population dynamics for stock assessments, which are used by resource managers to inform regulatory decisions. The large degree of individual variability we observed also suggests that random effects may be important to properly account for statistical dependencies arising from having repeated measures on the same individuals. Ignoring these dependencies will typically result in estimators of uncertainty that are biased low (Fieberg et al. 2010).

The simplest method for accommodating individual variation in habitat-selection studies is to fit separate models to each individual (Fieberg et al. 2010). Yet, a complication with this approach is that it becomes more challenging to estimate population parameters that quantify among-individual variability. For example, the variance of the individual coefficients will be impacted by both true variability and sampling error (Davidian 2017; Muff, Signer, and Fieberg 2020). Using the mixed ISSA, we can directly estimate these variance parameters and also use the same model for both individual and population-level inference. Furthermore, as our simulation example illustrates, we can obtain better estimates of individual-specific parameters by borrowing information from like individuals (Gelman and Hill 2006; Harrison et al. 2018). Yet, the application of mixed ISSAs can be challenging, especially when analyzing data sets containing small numbers of individuals. In our models, we included random effects for step length, log(step length) and cos(turn angle) but not their interactions with habitat class or distance to nearest edge habitat. This decision reduced the number of variance parameters that needed to be estimated but also constrained the degree of inter-individual variability that was included in our models. Complex models with many random effects can sometimes lead to computational issues, so these types of choices are commonplace when fitting generalized linear mixed effects models (Bolker et al. 2009). Additional guidance related to model-building strategies (e.g., methods for determining an appropriate level of model complexity) would be helpful. The two-step approach to parameter estimation (Avgar et al. 2016; Fieberg et al. 2021) and the Poisson trick for implementing mixed ISSAs (Muff, Signer, and Fieberg 2020) also complicates parameter interpretation and model inference (Michelot et al. 2023). Software implementations that directly fit models for mixed ISSAs in a single step would also be a welcome advance.

Individual variability has long been recognized as fundamental to understanding broad patterns in ecology, including those that emerge from the aggregation or interactions of smaller-scale individual units (Levin 1992). Recent attention has been given to methods that scale individual movement models to broader levels of animal space use or ecological dynamics. These studies include the use of step-selection analyses to derive utilization distributions and infer broad-scale patterns in space use (Potts and Börger 2023), and the integration of intra-specific variation from movement data in larger contexts such as community ecology (Costa-Pereira et al. 2022) and biodiversity studies (Jeltsch et al. 2013). Mixed ISSAs can be used to parameterize movement models for a population of individuals, which can then be simulated to explore population-level space-use patterns (Signer, Fieberg, and Avgar 2017; Signer et al. 2023). By providing annotated code with functions to easily fit and visualize models of animal movement using mixed ISSAs, we hope ecologists will take full advantage of their ability to quantify within- and among-individual variability in both movement and habitat-selection parameters to better understand ecological phenomena and species responses to changing environments.

## Declarations

All tagging protocols were approved by the Institutional Animal Care and Use Committee (# NCA19-002) of the North Carolina Aquariums. Research activities were carried out under a Scientific Research Permit issued to Nathan Bacheler by the Southeast Regional Office of the U.S. National Marine Fisheries Service, in accordance with the relevant guidelines and regulations on the ethical use of animals as experimental subjects. Mention of trade names or commercial companies is for identification purposes only and does not imply endorsement by the National Marine Fisheries Service, NOAA. The scientific results and conclusions, as well as any views and opinions expressed herein, are those of the authors and do not necessarily reflect those of any government agency.

## Authors’ contributions

NC led the analysis with contributions from DW, DK, JV, SF and JF; KS, JCT and NB collected the data; SF validated, documented, created figures and functions for implementing the analyses; JF, NC, and NB led the writing of the manuscript. All authors discussed, edited, read, and approved the final manuscript.

## Acknowledgements

We thank D. Hofmann for suggestions regarding the simulation. JF and NC were supported by National Aeronautics and Space Administration award 80NSSC21K1182 and JF received partial salary support from the Minnesota Agricultural Experimental Station.

## Appendix 1

### 8 Application of Mixed ISSAs to Snapper Data

This appendix provides additional details regarding the application of mixed ISSAs to the red snapper data. In particular, it includes a full set of equations and R code for specifying and summarizing the models.

#### 8.1 Integrated step-selection Analysis

Integrated step-selection analyses assume that animal space use is captured by the product of two functions, a movement-free habitat-selection kernel, *w*(*·*), that represents habitat preferences and a selection-free movement kernel, *ϕ*(*·*), that quantifies how the animal would move in the absence of habitat selection:

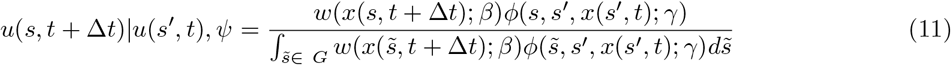

where *u*(*s, t* + Δ*t*) |*u*(*s*^*′*^, *t*), *ψ* represents the conditional probability of an animal being present at location *s* at time *t* + Δ*t*, given that it was at location *s*^*′*^ at time *t* and arrived there via the bearing *ψ*; *x*(*s*^*′*^, *t*) and *x*(*s, t* + Δ*t*) are habitat covariates measured at the start and end of the movement step, respectively; and *β* and *γ* are sets of parameters that quantify habitat selection and movement tendencies, respectively. The denominator in equation (1) integrates over the geographic area, *G*, that represents the area in which an animal could have potentially moved;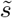 is a dummy variable for integration.

#### 8.2 Movement-Free Selection Function

In both mixed ISSAs, we modeled *w*(*·*) as a log-linear function of benthic habitat class and distance-to-edge measured at the end of the movement step:

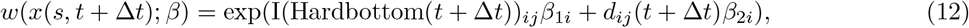

where *i* indexes the different individuals and *j* indexes the movement steps, *d*_*ij*_(*t* + Δ*t*) is the distance to edge habitat at the end of the movement step, and I(Hardbottom(*t* + Δ*t*))_*ij*_ is equal to 1 if the end of the movement step falls in the Hardbottom habitat class, and 0 otherwise. We used random effects to allow each individual to have its own set of habitat-selection parameters:

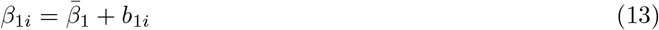

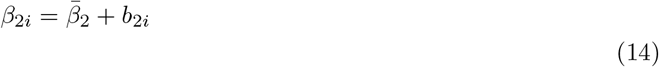

where 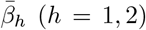 represents the mean coefficient across the population of individuals, and the *b*_*hi*_ are Normally distributed random effects associated with each explanatory variable in the model. We assumed the coefficients for Hardbottom habitat varied independently from the distance-to-edge coefficients:

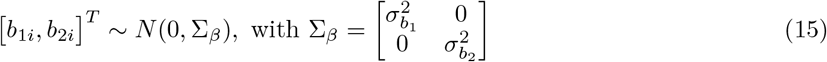

#### 8.3 Selection-Free Movement Kernels

##### 8.3.1 Model 1: Movement Dependent on Habitat Class

We assumed the step lengths associated with individual *i*’s movements starting in habitat class *k, sl*_*ik*_ followed a gamma distribution with individual-specific shape and scale parameters:

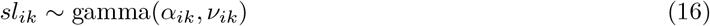

where (*k* = 1, 2) for sand and hardbottom, respectively. We assumed turn angles associated with individual *i*’s movements starting in habitat class *k, θ*_*ik*_ followed a von Mises distribution with individual-specific concentration parameters, *κ*_*ik*_, and with means, *µ*_*k*_ that were either 0 (assuming individuals had a tendency to move in a consistent direction) or ±*π* (assuming individuals were largely stationary or moving back and forth along linear features). We set *µ*_*k*_ to ±*π* and *κ* to |*κ*| when estimates of *κ* were negative and set *µ*_*k*_ 0 otherwise:

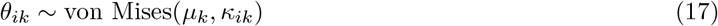

We included a single individual random effect for each type of parameter (*α, ν, κ*), which allowed the shape, scale, and concentration parameters to vary by individual, while forcing differences between habitat classes to be consistent across individuals.

To implement the model in R, we included step length, log(step length), and cos(turn angle) in the model along with interactions between these covariates and the benthic habitat class variable. We also included random effects for step length, log(step length), and cos(turn angle). We assumed the random effects were Normally distributed and allowed 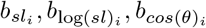 to co-vary independently from the habitat-selection coefficients.

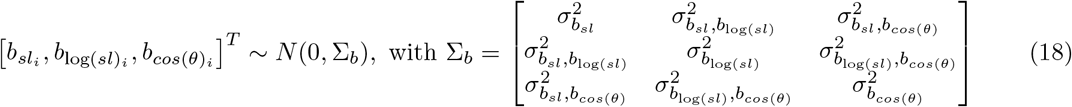

We can specify this model in R as follows:

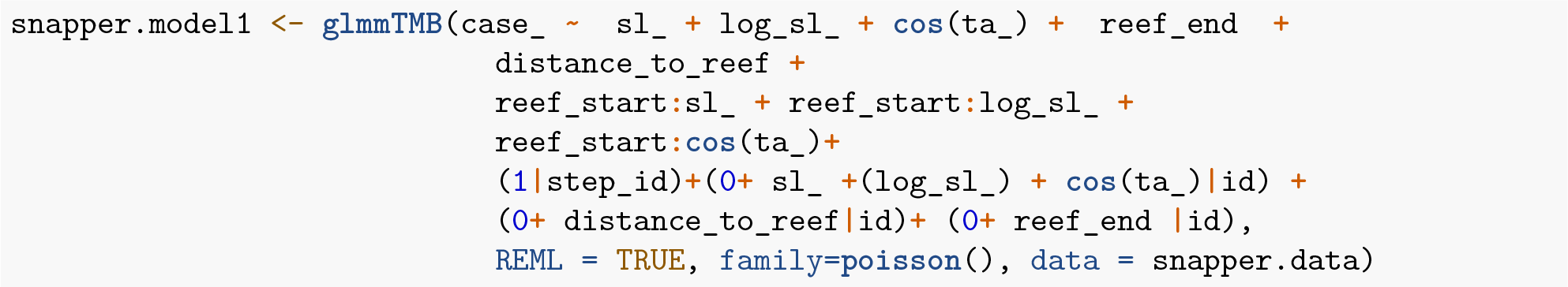

To connect the R specification to the formulas, note that:

1. `case_` is a binary variable equal to 1 for observed steps and 0 for random steps
2. `sl_, log_sl_` and `cos(ta_)` represent step lengths, log(step lengths), and cos(turn angle) for each step
3. `reef_end` and `distance_to_reef` measure the habitat class and distance to edge habitat at the end of the step (*I*(Hardbottom(*t* + Δ*t*))_*ij*_ and *d*_*ij*_(*t* + Δ*t*), respectively).
4. the interaction terms, `reef_start:sl_, reef_start:log_sl_` and `reef_start:cos(ta_)` are formulated in terms of the habitat class at the start of the movement step and allow the movement kernels to depend on the habitat class the fish is in.
5. `(1|step_id)` is included to implement the Poisson trick, making the model equivalent to a mixed conditional logistic regression (Muff, Signer, and Fieberg 2020).
6. `(0+ sl_ +(log_sl_) + cos(ta_)|id)` specifies that parameters of the gamma and von Mises distributions vary by individual and that these parameters have non-negative covariances.
7. `(0+ distance_to_reef|id) and (0+ reef_end |id)` specify that the habitat-selection parameters vary independently of each other and independently of the movement parameters.

#### 8.4 Obtaining updated movement parameters

Let *β*_*sl*_, *β*_log(*sl*)_, and *β*_*cos*(*θ*)_ represent the fixed-effects coefficients associated step length, log(step length) and cos(turn angle), respectively. Further, let 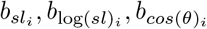 represent fish-specific random effects associated with these variables. In the first model, we estimate updated movement parameters for our reference class (sandy habitat) using:

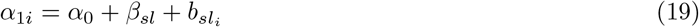

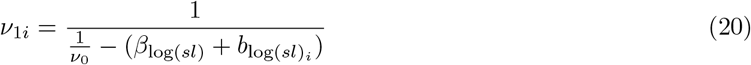

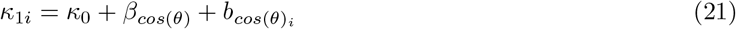

We can estimate updated parameters for the hardbottom habitat class by also including the coefficients for the interaction terms. For example, let *β*_*sl*:*I*(*Hardbottom*(*t*))_, *β*_*log*(*sl*):*I*(*Hardbottom*(*t*))_, and *β*_*cos*(*θ*):*I*(*Hardbottom*(*t*))_ be the interaction terms associated with the hardbottom habitat class at the start of the movement step. We can obtain updated estimates for *α*_2*i*_, *ν*_2*i*_, and *κ*_2*i*_ using:

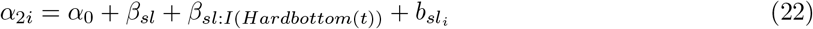

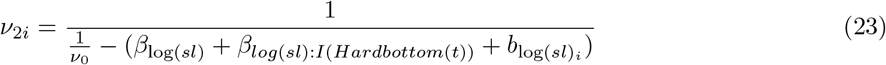

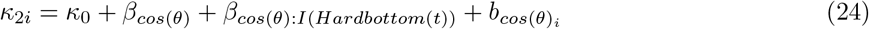

This update step can be accomplished in R using the mixedssf package as follows:

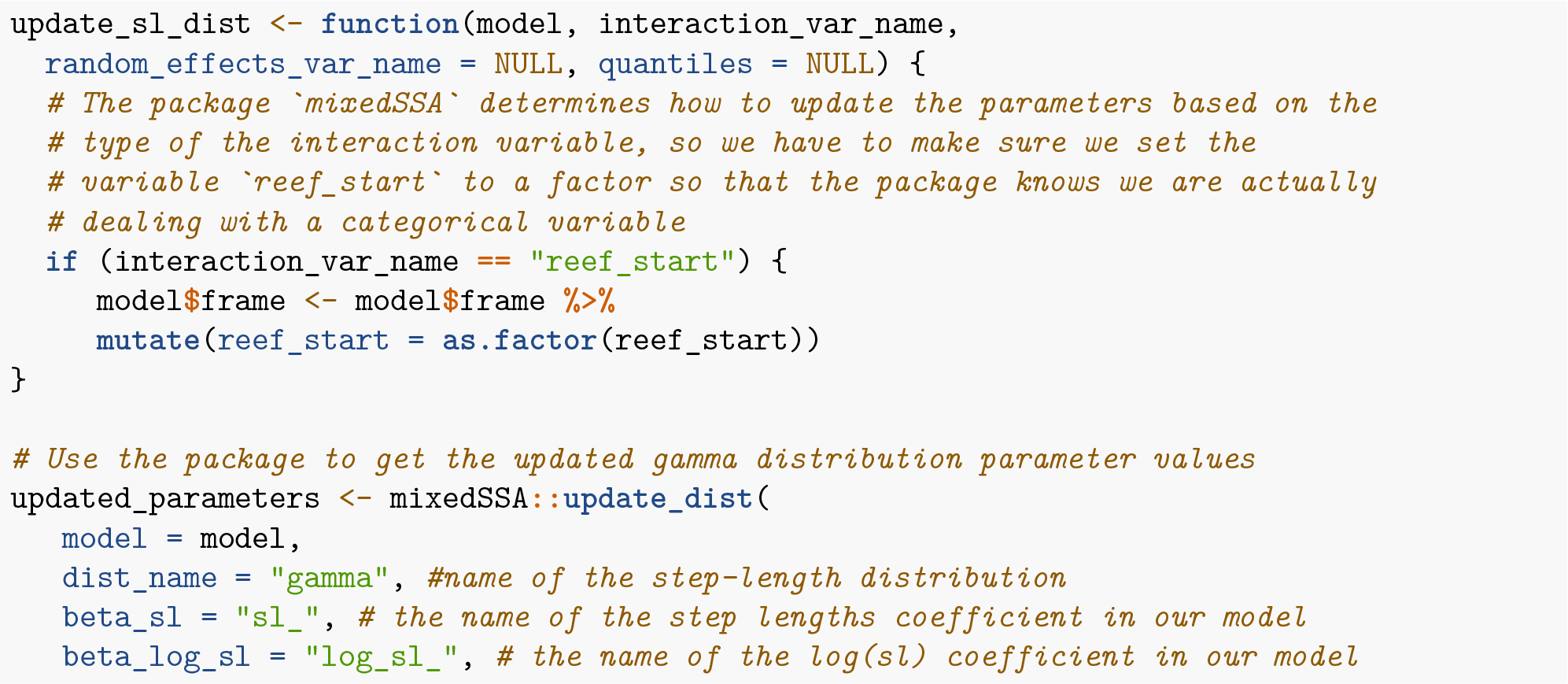
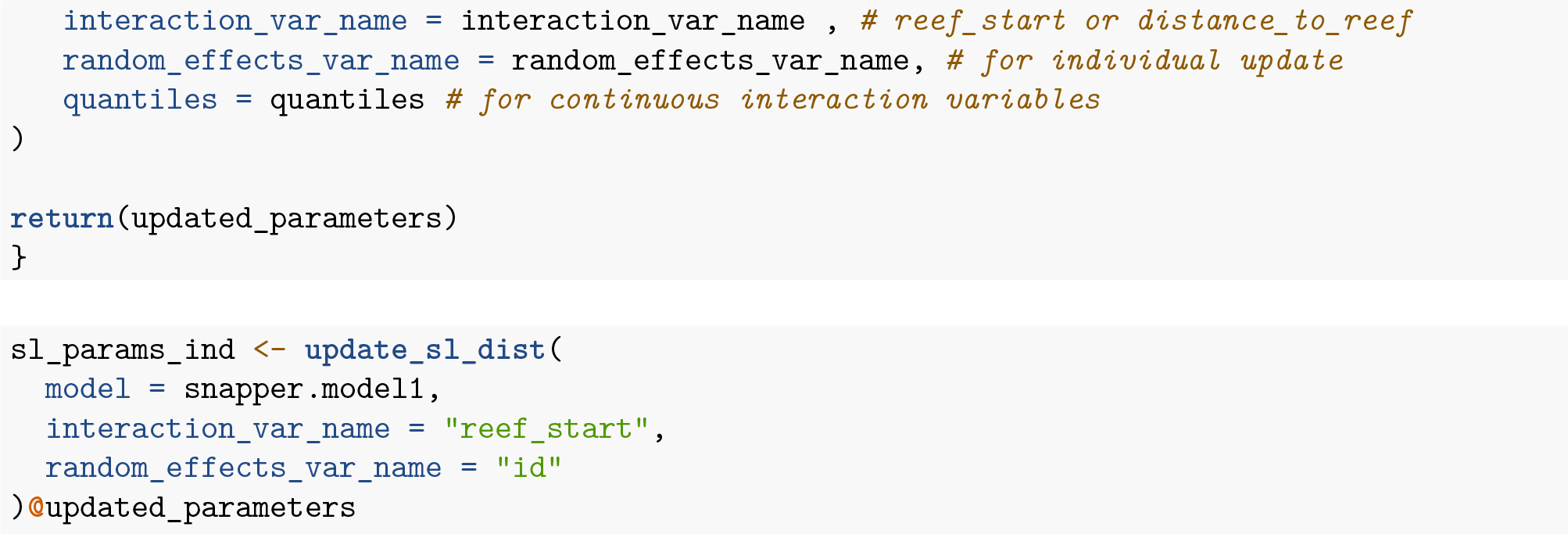

##### 8.4.1 Model 2: Movement Dependent on Distance to Reef Edge

We assumed the *j*^th^ step length for individual *i, sl*_*ij*_, followed a gamma distribution with individual-specific shape and scale parameters that depended on the distance from edge habitat, *d*_*ij*_(*t*), at the start of the movement step. We assumed turn angles, *θ*_*ij*_, followed a von Mises distribution with individual-specific concentration parameters, *κ*_*ij*_, that also depended on *d*_*ij*_(*t*). We used the same strategy for modeling *µ*_*ij*_ as with model 1 (setting *µ*_*ij*_ to 0 whenever *κ*_*ij*_ was positive and to ±*π* when *κ*_*ij*_ was negative).

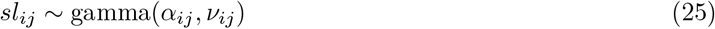

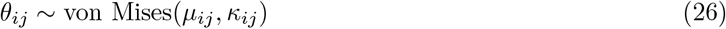

To implement the model in R, we again included step length, log(step length), and cos(turn angle) in the model along with interactions between these covariates and distance to edge habitat at the start of the step. We again included random effects for step length, log(step length), and cos(turn angle) but not the interactions with distance to edge habitat. The random effects were again assumed to be Normally distributed as in equation (18).

We can specify this model in R as follows:

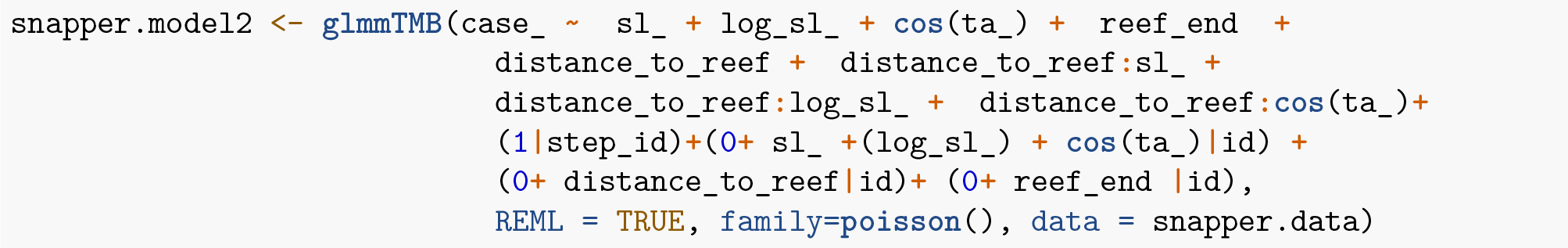

The terms in the model can be interpreted similarly to those in model 1.

##### 8.4.2 Updating the parameters of movement kernel

In our second model, our movement parameters were allowed to vary as a function of *d*_*ij*_(*t*). In that case, equations (19), (20), and (21) are used to estimate updated movement parameters when *d*_*ij*_(*t*) = 0. Let *β*_*sl*:*d*_, *β*_*log*(*sl*):*d*_, and *β*_*cos*(*θ*):*d*_ represent the interaction terms between step length, log(step length), cos(turn angle) and distance to the nearest edge habitat at the start of the movement step. We can obtain updated estimates of movement parameters for any other distance, *δ*, using:

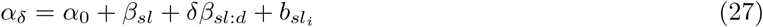

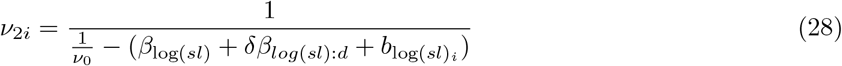

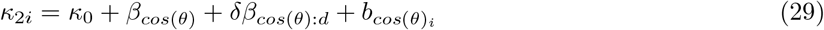

This update step can be accomplished in R using the mixedssf package as follows:

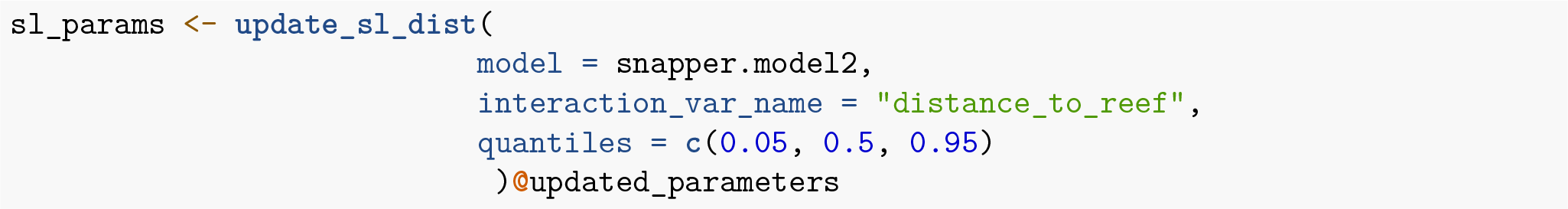

## Appendix 2

### 8.5 Supplementary Table

**Table 1:** Number of locations, start and end dates, and number of days tracked for individual fish in the study.

| id | Relocations | Start time | End time | No. of days tracked |
| --- | --- | --- | --- | --- |
| 2 | 18814 | 2019-05-07 08:32:37 | 2019-09-15 05:21:32 | 131 |
| 3 | 20785 | 2019-05-07 08:43:54 | 2019-08-28 10:42:55 | 113 |
| 4 | 604 | 2019-05-07 09:36:17 | 2019-05-10 18:58:42 | 3 |
| 5 | 15201 | 2019-05-07 08:53:12 | 2019-08-13 16:53:59 | 98 |
| 6 | 799 | 2019-05-07 08:57:33 | 2019-06-05 05:27:29 | 29 |
| 7 | 2755 | 2019-05-07 09:00:13 | 2019-06-26 17:22:27 | 50 |
| 8 | 16635 | 2019-05-07 09:19:17 | 2019-08-27 11:04:43 | 112 |
| 9 | 21297 | 2019-05-07 10:10:58 | 2019-12-16 07:51:58 | 223 |
| 10 | 4698 | 2019-05-07 10:33:04 | 2019-05-26 12:02:14 | 19 |
| 11 | 1290 | 2019-05-07 10:37:17 | 2019-06-19 09:57:00 | 43 |
| 12 | 22384 | 2019-05-07 10:56:51 | 2019-08-01 04:59:41 | 86 |
| 14 | 265 | 2019-05-07 11:06:47 | 2019-07-27 06:55:55 | 81 |
| 15 | 3005 | 2019-05-07 11:11:24 | 2019-05-20 17:51:28 | 13 |
| 16 | 52189 | 2019-05-07 11:19:54 | 2019-12-16 07:48:31 | 223 |
| 17 | 20316 | 2019-05-07 11:12:51 | 2019-07-29 07:58:04 | 83 |
| 18 | 5208 | 2019-05-07 11:30:55 | 2019-07-01 05:01:34 | 55 |
| 19 | 13505 | 2019-05-07 13:00:15 | 2019-07-14 05:40:40 | 68 |
| 20 | 12427 | 2019-05-07 12:53:10 | 2019-08-29 02:28:48 | 114 |
| 22 | 6755 | 2019-05-07 13:26:23 | 2019-06-20 03:29:39 | 44 |
| 23 | 1316 | 2019-05-07 14:14:46 | 2019-05-14 07:54:02 | 7 |
| 25 | 7979 | 2019-08-13 09:38:49 | 2019-11-11 12:04:01 | 90 |
| 26 | 1309 | 2019-08-13 10:16:40 | 2019-09-25 21:53:27 | 43 |
| 27 | 15277 | 2019-08-13 10:22:06 | 2019-12-15 16:15:58 | 124 |
| 29 | 4370 | 2019-08-13 10:30:32 | 2019-09-16 05:59:46 | 34 |
| 31 | 12774 | 2019-08-13 10:20:52 | 2019-11-09 09:57:45 | 88 |
| 32 | 7208 | 2019-08-13 11:00:50 | 2019-10-12 06:52:30 | 60 |
| 33 | 8586 | 2019-08-13 10:48:59 | 2019-09-30 06:55:07 | 48 |
| 37 | 22483 | 2019-08-13 12:30:40 | 2019-12-02 00:37:00 | 111 |
| 38 | 1766 | 2019-08-13 13:13:12 | 2019-12-15 16:14:22 | 124 |
| 39 | 3299 | 2019-08-13 12:49:03 | 2019-12-15 15:50:52 | 124 |
| 40 | 969 | 2019-08-13 12:54:02 | 2019-12-10 18:07:51 | 119 |
| 41 | 2324 | 2019-08-13 12:53:17 | 2019-12-15 16:18:55 | 124 |
| 42 | 450 | 2019-08-13 13:11:03 | 2019-11-27 06:28:52 | 106 |
| 46 | 10996 | 2019-08-30 11:39:53 | 2019-11-12 08:21:11 | 74 |
| 49 | 6253 | 2019-09-22 08:31:50 | 2019-11-06 04:51:00 | 45 |

**Table 2:** Coefficient estimates (95% CI) of the variables used in Model 1 and Model 2. Stars denote the significant p-value (+ p < 0.1, * p < 0.05, ** p < 0.01, *** p < 0.001).

| Variable | Model 1 | Model 2 |
| --- | --- | --- |
| sl_ <sup>a</sup> | -0.005 [-0.012, 0.002] | -0.007 [-0.015, 0.000]+ |
| log_sl_ <sup>b</sup> | 0.108 [0.042, 0.173]** | 0.072 [0.004, 0.140]* |
| cos(ta_) <sup>c</sup> | -0.855 [-0.927, -0.782]*** | -0.893 [-0.962, -0.823]*** |
| HardbottomEnd <sup>d</sup> | 0.247 [0.162, 0.332]*** | 0.235 [0.151, 0.318]*** |
| distance to edge | -0.013 [-0.021, -0.005]** | -0.023 [-0.032, -0.014]*** |
| sl_ X HardbottomStart <sup>e</sup> | -0.015 [-0.017, -0.012]*** | NA |
| log_sl_ × HardbottomStart | 0.053 [0.029, 0.076]*** | NA |
| cos(ta_) × HardbottomStart | 0.036 [-0.004, 0.076]+ | NA |
| sl_ × distance to reefedge | NA | 0.000 [0.000, 0.000]*** |
| log_sl_ × distance to reefedge | NA | 0.006 [0.005, 0.008]*** |
| cos(ta_) × distance to reefedge | NA | 0.006 [0.004, 0.007]*** |
| SD (sl_ id) | 0.02 | 0.021 |
| SD (log_sl_ id) | 0.175 | 0.181 |
| SD (cos(ta_) id) | 0.182 | 0.173 |
| SD (distance to reefedge id) | 0.021 | 0.023 |
| SD (HardbottomEnd id) | 0.208 | 0.203 |
| Cor (sl_~log_sl_ id) | -0.948 | -0.942 |
| Cor (sl_~cos(ta_) id) | 0.254 | 0.321 |
| Cor (log_sl_~cos(ta_) id) | -0.194 | -0.257 |
| Num.Obs. | 465488 | 465586 |
| R2 Marg. | 1 | 1 |
| R2 Cond. | NA | NA |
| AIC | 182377.9 | 182695.3 |
| BIC | 182576.8 | 182894.3 |
| RMSE | 0.22 | 0.22 |

**Note:** <sup>a</sup> Denotes length (m) of the step; <sup>b</sup> Denotes log(length) (m) of the step; <sup>c</sup> Denotes cosine of the turn-angle; <sup>d</sup> Equal 1 if the hardbottom benthic class at the end of the step; <sup>e</sup> Equal 1 if the hardbottom benthic class at the start of the step

**Table 3:** Estimated parameters in statistical distributions used to model step length and turn angles as a function of benthic habitat class (model 1) or distance to edge (model 2). The mean of the step-length distribution was calculated assuming a gamma distribution and the sd error was calculated using the delta method. The tentative distribution was observed from data and then movement parameters were updated using the coefficients of reef-class from model 1 and the distance to reef edge variable from model 2.

|  | shape | scale | mean | sd | kappa |
| --- | --- | --- | --- | --- | --- |
| Tentative | 1.090 | 18.240 | 19.884 | 1.382 | 0.348 |
| Sand | 1.198 | 16.794 | 20.115 | 1.305 | -0.506 |
| Hardbottom | 1.251 | 13.505 | 16.890 | 0.820 | -0.470 |
| 5%_quantile | 1.162 | 16.088 | 18.697 | 0.906 | -0.544 |
| 50%_quantile | 1.181 | 15.895 | 18.777 | 0.904 | -0.504 |
| 95%_quantile | 1.417 | 13.851 | 19.628 | 0.879 | -0.318 |

### 8.6 Supplementary figures

